# Widespread agrochemicals differentially affect zooplankton biomass and community structure

**DOI:** 10.1101/2020.10.01.322370

**Authors:** Marie-Pier Hébert, Vincent Fugère, Beatrix E. Beisner, Naíla Barbosa da Costa, Rowan D. H. Barrett, Graham Bell, B. Jesse Shapiro, Viviane Yargeau, Andrew Gonzalez, Gregor F. Fussmann

## Abstract

Anthropogenic environmental change is causing habitat deterioration at unprecedented rates in freshwater ecosystems. Despite increasing more rapidly than other agents of global change, synthetic chemical pollution –including agrochemicals such as pesticides– has received relatively little attention in freshwater biotic assessments. Determining the effects of multiple agrochemicals on complex community and ecosystem properties remains a major challenge, requiring a cross-field integration of ecology and ecotoxicology. Using a large-scale array of experimental ponds, we investigated the response of zooplankton community properties (biomass, composition, diversity metrics) to the individual and joint presence of three widespread agrochemicals: the herbicide glyphosate, the neonicotinoid insecticide imidacloprid, and fertilisers. We tracked temporal variation in community biomass and structure (i.e., composition, diversity metrics) along single and combined pesticide gradients (each spanning eight levels), under low (mesotrophic) and high (eutrophic) nutrient-enriched conditions, and quantified (i) agrochemical interactions, (ii) response threshold concentrations, and (iii) community resistance and recovery. We found that major zooplankton groups differed in their sensitivity to pesticides: ≥3 µg/L imidacloprid impaired copepods, rotifers collapsed at glyphosate levels ≥0.3 mg/L, whereas some cladocerans were highly tolerant to pesticide contamination. Glyphosate was the most influential driver of community properties, with biomass and community structure responding rapidly but recovering unequally over time. Zooplankton biomass showed little resistance when first exposed to glyphosate, but rapidly recovered and even increased with glyphosate concentration; in contrast, richness declined in more contaminated ponds but failed to recover. Our results show that the biomass of tolerant taxa compensated for the loss of sensitive species, conferring greater resistance upon subsequent exposure; a rare example of pollution-induced community tolerance in freshwater metazoans. Overall, zooplankton biomass appears to be more resilient to agrochemical pollution than community structure, yet all community properties measured in this study were affected at glyphosate levels below common water quality guidelines in North America.

## 1. Introduction

Freshwater ecosystems have been extensively altered by human-induced global change and environmental degradation. A critical, yet largely understudied dimension of global change is synthetic chemical pollution (Mazor et al., 2018; Rockström et al., 2009; Steffen et al., 2015), including the release of agrochemical contaminants such as pesticides in ecosystems (Malaj et al., 2014; Schäfer et al., 2016). Pesticide manufacturing has risen massively since agricultural industrialisation, expanding more rapidly than other well-recognised anthropogenic drivers of global change (Bernhardt et al., 2017). With a global annual application of pesticides exceeding four million tonnes (FAOSTAT 2020; Maggi et al., 2019), their presence in surface and ground waters is increasingly reported (Ippolito et al., 2015; Stehle & Schulz, 2015; Wittmer et al., 2010), in addition to the recurrent presence of other agrochemicals such as fertilisers (Vörösmarty et al., 2010). Despite these global trends, few studies addressed how the rising load and diversity of synthetic agrochemicals entering freshwaters affect the structure and function of communities and ecosystems (Gessner & Tlili, 2016; Mazor et al., 2018; Reid et al., 2019).

Ecotoxicological assessments of agrochemical effects on freshwater biota often rely on laboratory assays conducted in simplified and controlled settings, facilitating the identification of modes of action and threshold concentrations causing impairment. Although crucial to establishing baseline knowledge of synthetic chemicals, laboratory toxicity tests typically focus on single species and agrochemicals, ignoring the possible influence of many indirect and interacting factors on multispecies assemblages (Fleeger et al., 2003; Rohr et al., 2006). Long-standing needs to better transpose observations from ecotoxicological studies to our understanding and management of ecosystems include several key considerations: (i) a broader appreciation of how species and trophic interactions may modulate responses to pollutants (Relyea & Hoverman, 2006; 2008); (ii) determining interactions of co-occurring agrochemicals (Coors & De Meester, 2008; Relyea, 2009); (iii) incorporation of temporal dynamics to assess time-dependent effects of agrochemicals or community processes and recovery (Halstead et al., 2014; Rohr & Crumrine, 2005); (iv) accounting for physicochemical factors (e.g., exposure to natural light; Fenoll et al., 2015) that may influence the flow, toxicity, and degradation of agrochemicals, especially pesticides. Field experiments –such as whole-ecosystem manipulations or outdoor mesocosms– are complementary in this regard, as they can demonstrate how the structure and function of biological systems may be influenced under realistic conditions, expanding the scale of inquiry in ecotoxicology and tackling processes operating at higher levels of organisation (communities, ecosystems; Rohr et al., 2006; Peters et al., 2013; Schmitt-Jansen et al., 2008). Echoing earlier calls, recent studies stressed the ever-critical need to better integrate ecology and ecotoxicology to develop effective risk assessments and conservation strategies adapted to complex environments confronted with rising agrochemical pollution (Bernhardt et al., 2017; Gessner & Tlili, 2016).

A diversity of pesticides and fertilisers co-occur in surface waters, in part because it is common practice to apply agrochemicals as mixtures (Altenburger et al., 2013). Within the U.S. alone, traces of pesticides are nearly ubiquitous in lotic systems, with >90% of the streams located in agricultural, urban, and mixed land use areas having detectable levels of at least two pesticides (Gilliom et al., 2006). The concomitant presence of agrochemicals can generate interactive effects that may weaken or strengthen individual effects of agrochemicals on biota, resulting in nonadditive outcomes (synergies or antagonisms) that are hard to predict from single agrochemical studies (Geyer et al., 2016; Relyea, 2009). Previous assessments of multiple stressor effects on freshwater biota reveal that cumulative effects are more often interactive than additive (Birk et al., 2020; Jackson et al., 2016). For example, Chará-Serna et al. (2019) found that imidacloprid, a neonicotinoid insecticide, mitigated the positive effect of nutrients on freshwater invertebrate richness. The study of multiple stressors, however, has thus far primarily focused on characterising interaction types across organisms, not along stress gradients; in fact, the paucity of regression-style designs has been recognised as a hindrance to assess effect sizes and potential response thresholds (Kreyling et al., 2018; Orr et al., 2020). Although stressor interactions represent a key issue for aquatic conservation (Côté et al., 2016; Reid et al., 2019), interactions of pesticides remain rarely addressed, with syntheses reporting few studies of synthetic chemicals and xenobiotics other than fertilisers (Birk et al., 2020; Jackson et al., 2016).

Adding further complexity to the study of agrochemicals is that different community properties may show distinct responses to multiple pollutants. Examining temporal variation in aggregate community properties –such as standing biomass, species assemblages, or particular functions– aids in identifying the underlying processes allowing communities to resist or recover from stressors (Halstead et al., 2014; Hillebrand et al., 2018; Kéfi et al., 2019). Faced with a perturbation, community properties may show resistance (i.e., ability to remain unaffected upon disturbance) or recovery (i.e., ability to return to initial state after disturbance). In some cases, the resistance or recovery of a given property can be achieved via the lack of resistance or recovery in another property; a trade-off that has been observed between community structure (i.e., composition, diversity) and biomass. For instance, community structure may rearrange under stress (e.g., turnover or loss of taxa; Murphy & Romanuk, 2014), enabling biomass stocks and production to be maintained, and thus preserving a key ecosystem function (Allan et al., 2011; Gonzalez & Loreau, 2009; Hoover et al., 2014). In such scenario, species sorting results in the resistance of community biomass, implying a relatively weaker resistance of community structure. This trade-off, whereby biomass resistance is accomplished at the expense of community structure, may occur if taxa that are tolerant to multiple stressors compensate for more sensitive community members; a process that may be influenced by the diversity and co-tolerance patterns within the original species pool (Arnoldi et al., 2019; Cottingham et al., 2001; Vinebrook et al., 2004). Analogous trade-offs can also occur between the recovery of community structure and biomass over time (Hillebrand & Kunze, 2020). Furthermore, if stress-induced species sorting results in the replacement of sensitive taxa with tolerant ones, community-wide tolerance may increase, leading to greater resistance upon subsequent stress exposure (i.e., stress- or pollution-induced community tolerance; Tlili et al., 2016; Vinebrook et al., 2004). However, if too few taxa are tolerant, communities may collapse entirely, leading to cascading food web and ecosystem effects (Dunne & Williams, 2009), such as secondary extinctions (Ives & Cardinale, 2004). For example, increased use of neonicotinoid insecticides in agricultural watersheds can induce a decline in zooplankton and the subsequent collapse in the yield of a fishery (Yamamuro et al., 2019). Overall, assessing resistance and recovery of aggregate community properties may reveal mechanisms by which biota cope with stress, providing in the context of this study a clearer picture of immediate versus long-lasting effects of agrochemical pollution.

In this study, we address how complex zooplankton communities respond to the individual and cumulative effects of fertilisers (nutrient pollution) and two widely used, globally relevant pesticides: the herbicide glyphosate and the neonicotinoid insecticide imidacloprid. Glyphosate- and imidacloprid-based pesticides are extensively used in agricultural landscapes (Maggi et al., 2020; Simon-Delso et al., 2015) and can reach aquatic ecosystems in different ways (Hébert et al., 2019; Jeschke et al., 2011; Medalie et al., 2020; Struger et al., 2017), leading to their widespread occurrence in freshwaters (Aparicio et al., 2013; Montiel-Léon et al., 2019; Morrissey et al., 2015) and concerns over their toxicity to aquatic life (Anderson et al., 2015; van Bruggen et al., 2018). Zooplankton have been extensively used as model organisms in toxicological assessments on non-target aquatic biota. Table S1 provides a non-exhaustive compilation of (>40) experimental studies addressing the effects of glyphosate- and imidacloprid-based pesticides on freshwater zooplankton; reflecting the long-standing contrast between the many short-term, single-species laboratory tests and the relatively scarce field-based evaluations of communities exposed to multiple agrochemicals. Although extensive laboratory research has demonstrated the adverse effects of glyphosate and imidacloprid, these may be undetectable (or reversed, i.e., positive influence) through other testing approaches (Table S1; Mikó et al., 2015). The wide range of sensitivity across zooplankton taxa, as well as the discrepancy between model species used in laboratory tests (e.g., *Daphna magna*) and those found to be responsive in natural assemblages further highlight the need to examine how these pesticides may affect complex communities in polluted freshwater environments.

Using a large outdoor array of 48 experimental ponds, we performed a 43-day study to track temporal variation in zooplankton (crustacean and rotifer) community properties (biomass, composition, and three diversity metrics) along single and combined gradients of glyphosate and imidacloprid, under low (mesotrophic) and high (eutrophic) nutrient-enriched conditions. We contrasted responses of zooplankton biomass and community structure (hereby measured via species composition and diversity metrics), and specifically quantified (i) interactive effects of agrochemicals, (ii) response threshold concentrations, and (iii) resistance and recovery patterns across community properties. For each objective, we formulated general predictions in Table 1. Our approach is aligned with recent calls to bridge ecotoxicology and ecology, addressing potential interactions of multiple co-occurring agrochemicals along gradients of stress, with respect to response thresholds. Our results have implications for freshwater zooplankton exposed to globally relevant agricultural pollutants at levels both below and above common North American water quality guidelines. Finally, through the consideration of community resistance and recovery across multiple aggregate properties, we position our findings within the broader field of ecological stability and ecosystem responses to global change.

**TABLE 1.** Summary of initial predictions for each objective.

| **Treatment** | **Prediction** |
| --- | --- |
| *General effect of agrochemicals* |  |
| Glyphosate | Negative effect on zooplankton (based on Table S1) |
| Imidacloprid | Negative effect on zooplankton (based on Table S1) |
| Nutrient fertilisers | Positive (resource-mediated) effect on zooplankton |
| *Agrochemical additive versus interactive effects* |  |
| Combination: glyphosate, nutrients | N will mitigate the negative effects of G;<br>possible positive interaction |
| Combination: imidacloprid, nutrients | N will mitigate the negative effects of I;<br>possible positive interaction |
| Combination: glyphosate, imidacloprid | G and I will result in greater negative effects;<br>possible negative interaction |
| Combination: glyphosate, imidacloprid, nutrients | N will mitigate the combined negative effects of G and I;<br>possible positive or negative interaction |
| *Response thresholds* |  |
| Glyphosate | Negative effects observed at 0.2 mg/L or higher (≥ dose 4)<br>(based on Table S1) |
| Imidacloprid | Negative effects observed at 0.3 mg/L or higher (≥ dose 3)<br>(based on Table S1) |
| *Community resistance and recovery* |  |
| Single or joint presence of pesticides | Biomass will show greater resistance and recovery than<br>community structure (composition and diversity metrics) |

G = glyphosate (herbicide); I = imidacloprid (insecticide); N = nutrient fertilisers

## 2. Methods

### 2.1 Experimental design and treatments

We conducted a field experiment at the Large Experimental Array of Ponds (LEAP), a pond mesocosm facility built at McGill University’s Gault Nature Reserve (Mont St-Hilaire, Québec, Canada), a protected, forested area. LEAP is connected via a 1-km pipe to the headwater Lake Hertel from which water and organisms can flow by gravity and accumulate in a large reservoir; once mixed, water and organisms can be evenly distributed across 100 experimental ponds, each having a capacity of *ca*. 1,000L.

On 11 May 2016, we filled all experimental ponds with lake water and planktonic organisms, followed by a three-month acclimation period. We removed fish using a coarse sieve when filling ponds, and periodically removed tadpoles and debris with a net. Every two weeks, we replaced 10% of the pond volume with a fresh lake inoculum to track seasonal changes in Lake Hertel’s plankton community, maximise the diversity of the initial pool of species, and minimise ecological drift across ponds prior to the experiment. We recorded physicochemical variables and water level weekly to track homogeneity across ponds. On 10 August 2016, we selected 48 ponds for a collaborative experiment to assess the responses of planktonic communities, including phytoplankton and bacterioplankton (see Barbosa da Costa et al., 2020; Fugère et al., 2020). Here we only report observations made over the first 43 days, as zooplankton communities were no longer sampled at the same resolution beyond this time point.

Agrochemical treatments consisted of two nutrient-enriched levels, mimicking mesotrophic (ambient Lake Hertel state) and eutrophic conditions, and three pesticide gradients spanning eight levels: glyphosate alone, imidacloprid alone, and a combination of both pesticides; see schematic representation in Figure 1a. A regression design for pesticide application enabled the quantification of effect strengths while identifying threshold concentrations affecting zooplankton. Target doses of glyphosate (acid equivalent) were 0 (control), 0.04, 0.10, 0.30, 0.70, 2.00, 5.50, and 15.0 mg/L; while imidacloprid were 0 (control), 0.15, 0.40, 1.00, 3.00, 8.00, 22.0, and 60.0 µg/L (Figure 1b). Pesticide levels covered a range of concentrations in line with those from ecotoxicological studies (Table S1), while spanning benchmarks considered safe for aquatic life (Canadian Water Quality Guidelines; CCME 2007; 2012; Figure 1b). Target concentrations were set to maintain a constant (logarithmic) increment across doses. For glyphosate, none of the doses exceeded short-term criteria for aquatic life in Canada (CCME 2012; Figure 1b) or for freshwater invertebrates in the United States (Office of Pesticide Program; EPA 2019).

**FIGURE 1.**
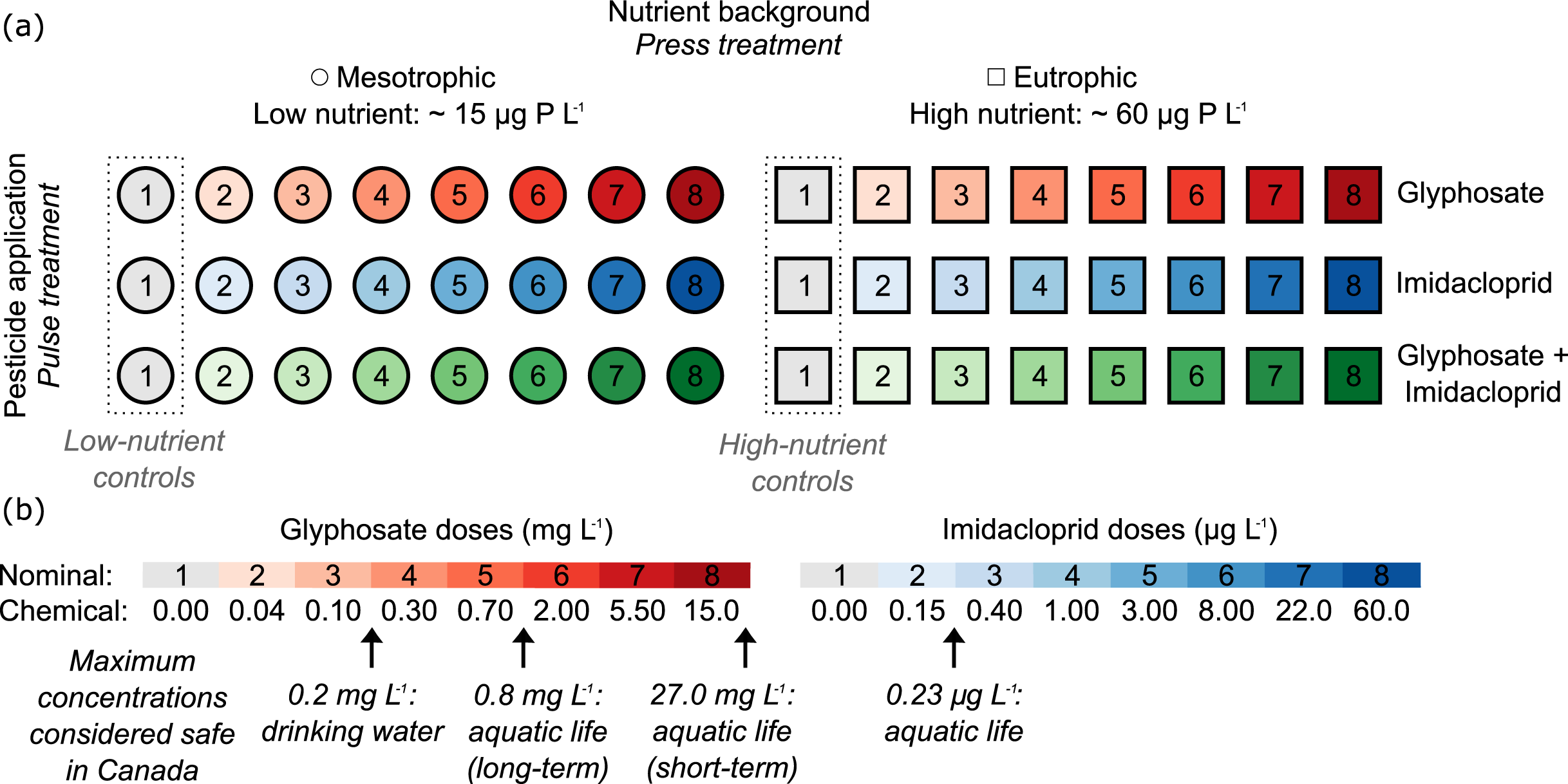
Simplified scheme of the experimental design. (a) Nutrient treatment (two levels) and pesticide gradients (8 doses for each of the three gradients) applied to the 48 experimental ponds. Nutrients were applied as a press treatment, whereas pesticides were applied twice in the form of pulses on days 6 and 34. Symbols correspond to low- and high-nutrient treatments. Colour type indicates pesticide treatment type (red = glyphosate only, blue = imidacloprid only, green = both pesticides); colour saturation and numbers refer to target pesticide doses upon the application of the first pulse. Control ponds, wherein only nutrient background was manipulated, are denoted in gray. (b) Nominal and chemical doses of pesticides are represented with respect to water quality threshold concentrations in Canada (CCME 2007; 2012).

To mimic the natural flow of agrochemicals to surface waters, we applied nutrients as a press treatment, and pesticides in the form of two pulses. We manipulated nutrient enrichment while maintaining the same nitrogen (N) to phosphorus (P) ratio as our source lake (∼33). Target P levels were 15 µg P/L (mesotrophic; referred to as low nutrient) and 60 µg P/L (eutrophic; high nutrient); we first added nutrient solutions (prepared with KNO3, KH_2_PO_4_, and K_2_HPO_4_) on 10 August 2016. To maintain a press treatment, this step was repeated every two weeks. We started our 43-day experiment on 16 August (day 1). On days 6 and 34, we applied pesticide treatments. We prepared glyphosate solutions with Roundup Super Concentrate (Monsanto©); calculations to reach target concentrations were based on the glyphosate acid content of the formulation. We prepared imidacloprid-based solutions by dissolving imidacloprid powder (Sigma-Aldrich) in ultrapure water.

### 2.2 Sampling

We sampled all ponds on six occasions: days 1, 7, 15, 30, 35, and 43. Our sampling schedule included one instance prior to the first pesticide pulse on day 6, three time points between the two pulses, and two after the second pulse on day 34. The relatively long interval between the two pulses (28 days) was intended to permit the potential recovery of communities prior to the second pulse. On each sampling day, we collected water with 35-cm long integrated tube samplers at multiple locations within ponds and stored samples in dark Nalgene© bottles for nutrient and biotic measurements other than zooplankton. To avoid cross-contamination, we assigned each pond its own sampler and bottles. We immediately transferred samples to our on-site indoor laboratory and kept in the dark at 4°C. For zooplankton, we collected water at multiple locations in each pond using our integrated samplers and sieved 2L with a 64-µm mesh. We anesthetised zooplankton with carbonated water and samples were fixed with ethanol (final concentration ∼ 75%) on site. We measured a standard suite of physicochemical parameters using a YSI© multiparameter sonde. To measure pesticides concentrations, we collected samples outside of our regular sampling schedule, as pesticides are subject to partial degradation in water. We sampled ponds immediately after pesticide application, on days 6 and 34; we collected additional samples on days 14 and 29 in a subset of the ponds to track degradation over time. We acidified pesticide samples (pH below 3) and kept them frozen at -20°C until analysis.

### 2.3 Laboratory analyses

We carried out a thorough analysis of zooplankton community composition for all 48 experimental ponds of the six sampling occasions (N=288). For the focal 288 samples of this study, we counted and identified every crustacean and rotifer individual (spanning 24 taxa) using an Olympus dissecting scope and an Olympus inverted microscope.

We analysed nutrient samples in the GRIL laboratory at the University of Québec at Montréal. We oxidised TP and TN samples with persulfate and alkaline persulfate digestions, respectively. We measured TP as orthophosphate using the spectrophotometric molybdenum method (890nm; Ultrospec 2100 pro, Biochrom), and TN as nitrites (reduced via a cadmium reactor) with a flow analyser (OI-Analytical Flow Solution 3100). We measured pesticides concentrations via liquid chromatography coupled to mass spectrometry (LC-HRMS) using an Accela 600-Orbitrap LTQ XL (LC–HRMS, Thermo Scientific) in the Department of Chemical Engineering at McGill University. The methods used are more extensively described in Fugère et al. (2020; glyphosate) and Barbosa da Costa et al. (2020; imidacloprid). Briefly, we measured glyphosate via heated electrospray ionization in negative mode (mass range = 50–300 m/z), and imidacloprid in positive mode (mass range = 50–700 m/z). We used an ion trap to perform targeted data acquisition for the product ion spectra (MS2) and generate identification fragments.

### 2.4 Data manipulation and statistical analyses

To enable comparison across crustaceans and rotifers, we converted abundance data to biomass using taxon-specific individual dry mass estimates compiled by Gsell et al. (2016) and Hébert et al. (2016). We then applied a log_10_(1+X)-transformation on biomass data. In analyses, we treated nutrient treatment as a binary factor variable, and used nominal levels of treatment doses for pesticides. We performed all statistical analyses in R version 4.0.0 (R Development Core Team).

We explored relationships with physicochemical factors but included none as predictors in models as there was little variation across ponds. We also examined the potential role of chlorophyll-*a* (chl-*a*) as a driver of zooplankton biomass. However, chl-*a* was highly correlated with glyphosate and TP after pulse 1 (presumably due to fertilising effects of glyphosate-derived P; see Discussion). To avoid the inclusion of highly correlated predictors, we decided against including chl-*a* in our main models. The rationale behind this decision was that our ultimate goal was to assess the causal, potentially cascading effects of agrochemicals on zooplankton, not to disentangle direct from indirect effects (e.g., mediated via changes in chl-*a*). As such, regardless of whether zooplankton biomass increased with chl-*a* as a result of glyphosate fertilising effects over time, this indirect effect was triggered by the presence of glyphosate in water, which is ultimately the effect that we aimed to quantify. We nonetheless present relationships between chl-*a* and zooplankton biomass in the Supplementary Material section (Table S4; Figure S3).

#### Effects on community biomass

We quantified the effects of time, agrochemicals, and all interactions on total and group-specific zooplankton biomass, using linear mixed effect models (LMM). To account for pseudoreplication (non-independence) across temporally repeated measurements from the same ponds, we set “individual pond” as a random effect. Predictors were standardised as per Gelman (2008; i.e., centered and divided by two standard deviations) prior to running LMMs so as to adequately compare effect sizes between continuous (pesticide dose) and binary (nutrient level) predictors. To fit LMMs, we used the function *lmer* in the R package *lme4*. We report model marginal R^2^ and conditional R^2^, representing proportions of variance explained by fixed factors alone (i.e., treatment and time) and both fixed and random factors, respectively. We also report intraclass correlation coefficients (ICC) as an estimate of within-pond correlation across temporally repeated measures.

Given the time-dependence of effects (i.e., direction of effect reversing over time), we quantified the effects of each agrochemical (nutrients, glyphosate, imidacloprid) and their interactions (nutrients:glyphosate, nutrients:imidacloprid, glyphosate:imidacloprid) for each sampling day, fitting all effects as interaction effects with time (converted to a factor). We also tested higher-order interactions but in none of the model were they significant; thus, they are not presented. We quantified parameter estimates for day 1 (Tables S3-S7) but excluded them from main figures, as pesticides were only applied as of day 6. We validated LMMs through the examination of residuals (distribution and homoscedasticity). We graphically represented effect sizes (measured as model parameter estimates) in forest plots using the *sjplot* and *ggplot2* packages in R.

To identify threshold concentrations affecting zooplankton, we used univariate regression trees (URT). Through recursive partitioning, URTs repeatedly divide data to identify predictor values associated with abrupt changes in response data; predictor values are retained as ‘thresholds’ (breaking points or splits) when data are divided such that the sums of squares of the groups created by the tree are minimised. URTs provide complementary information to LMMs, as the former can detect non-linear effects and specifically identify doses causing biotic responses (unlike LMMs that quantify overall effects across sites/doses). We included nominal pesticide levels, nutrient status, and time as predictors in URTs, and used the *ctree* function in the R package *party*. To preclude overfitting, we restricted models to a maximum of 4 splits, only allowed when *P* < 0.01; *P*-values were estimated by permutation tests as per Hothorn et al. (2006).

#### Effects on community structure: composition and diversity metrics

To assess how community structure varied across ponds and over time, we measured changes in taxonomic composition and diversity metrics, using a series of multivariate (ordinations; composition) and univariate (diversity metrics) analyses.

We discarded data from two low-nutrient, glyphosate-treated sites (doses 7 and 8) from day 7, as those samples did not contain any zooplankton (temporary collapses), making them unfit for compositional analyses. We log-transformed biomass data as per Anderson et al. (2006) to reduce asymmetry. To visualise compositional changes over time with respect to treatment, we used principal component analysis (PCA). We built PCAs using the *rda* function of the R package *vegan* (Oksanen et al., 2019). To better illustrate temporal dynamics, we constructed the multidimensional space of the ordination using pond data from all sampling days; then, we represented time-specific compositional data in six different panels. By doing so, we incorporated temporal signals in PCA scores. We then used PCA scores to: (1) quantify the effect of treatments and time on community composition using a LMM, built with the same structure as the biomass models described previously; and (2) identify breaking points in compositional shifts using URTs.

To more clearly visualise temporal shifts in species assemblages, and identify which taxa were most responsive to treatment and thus responsible for compositional changes, we used principal response curves (PRC; Van den Brink et al., 1999; 2008). PRCs are a type of redundancy analyses contrasting divergence in composition between reference (control) and perturbed (treated) sites in a chronological fashion. The graphical output of PRCs illustrate the degree to which treated communities deviate (left Y axis) from controls (horizontal line where Y=0) over time (X axis). We averaged replicates of control communities based on their nutrient treatment (see Figure 1a) to quantify a mean community matrix for each nutrient level. Using the *prc* function in the R package *vegan* (Oksanen et al., 2019), we built six PRCs for each agrochemical combination, and an additional one using only low- and high-nutrient control ponds to quantify the effect of nutrient enrichment alone. Using the first constrained axis, we examined the proportion of the variance explained by (i) time alone (conditional coefficient), and (ii) the interaction between treatment and time (constrained coefficient). Species scores are projected on the right Y axis as a taxon-specific measure of responsiveness to treatment and contribution to overall compositional changes. Score signs indicate the direction of response (positive: increases in density; negative: decreases in density) relative to control communities; only scores higher than 0.5 (in absolute value) were retained as significantly responsive.

To assess how agrochemicals affected community diversity, we examined variation in alpha diversity (exponent of the Shannon index), richness (taxon number), and evenness (Pielou’s index). We calculated diversity metrics for crustaceans, rotifers, and the whole zooplankton community, left metrics untransformed in analyses. We quantified effects of treatments using LMMs and URTs, consistently with biomass and composition models.

#### Resistance and recovery measures

Using the framework developed in Hillebrand et al. (2018), we explored resistance and recovery of four community properties (biomass, composition, richness, and alpha diversity), with the aim of comparing responses between biomass and community structure (i.e., composition and diversity metrics). We hereby define community resistance as the ability of a community to withstand a perturbation (i.e., similarity in community properties between control and treated sites immediately after perturbation), and recovery as the ability to return to the state in which the community would be in the absence of a perturbation (i.e., similarity in community properties between control and treated sites some time after perturbation). For each pesticide dose, we estimated resistance to treatment using measurements made on sampling days 7 and 35 (i.e., 24-36 hours after pesticide application), and used data from subsequent days (15, 30, 43) to track recovery after pulses. We averaged replicates of control communities based on their nutrient level (consistently with our approach to estimate a mean control community in PRCs). To quantify the resistance and recovery of community properties, we used effect size measures between controls and pesticide-treated ponds as per Hillebrand et al. (2018). For biomass, richness, and diversity, we calculated the log response ratio (LRR) between controls and treated communities. In this framework, LRR=0 indicates full resistance or recovery; LRR<0 indicates low resistance or incomplete recovery; and LRR>0 indicates low resistance or recovery via overcompensation/stimulation. For composition, we calculated the similarity between controls and treated ponds using the Bray-Curtis dissimilarity index, resulting in values ranging from 0 and 1, whereby 1 indicates full resistance/recovery.

For each pesticide pulse, we examined resistance and recovery trends across community properties via three types of relationships: (i) between (resistance/recovery) measures of community properties and pesticide dose (Figure S7), (ii) between (resistance/recovery) measures of two different community properties (Figures 7, S8), and (iii) between resistance and recovery within community properties (Figure S9). We used Spearman nonparametric rank correlation coefficients to assess the strength of relationships. As mentioned, two zooplankton samples were empty, resulting in LRRs of -∞ for biomass and richness; we graphically illustrated those measures (identified via gray layers) but excluded them from the calculation of correlation coefficients.

## 3. Results

### 3.1 Effects of agrochemicals on biomass

Over the 43 days of this experiment, the biomass of zooplankton communities ranged from 0 to 1667.3 µg/L (dry mass; 0–1142.5 organisms/L), with a mean of 128.8 µg/L (98.3 organisms/L). On day 1, one week prior to the first pulse of pesticides, zooplankton biomass did not differ between low- and high-nutrient ponds (Figure 2). A LMM revealed that nutrient enrichment alone did not affect overall zooplankton biomass throughout the experiment, nor did it affect the biomass of major taxonomic groups (i.e., cladocerans, copepods, and rotifers; Figure 3a-d).

**FIGURE 2.**
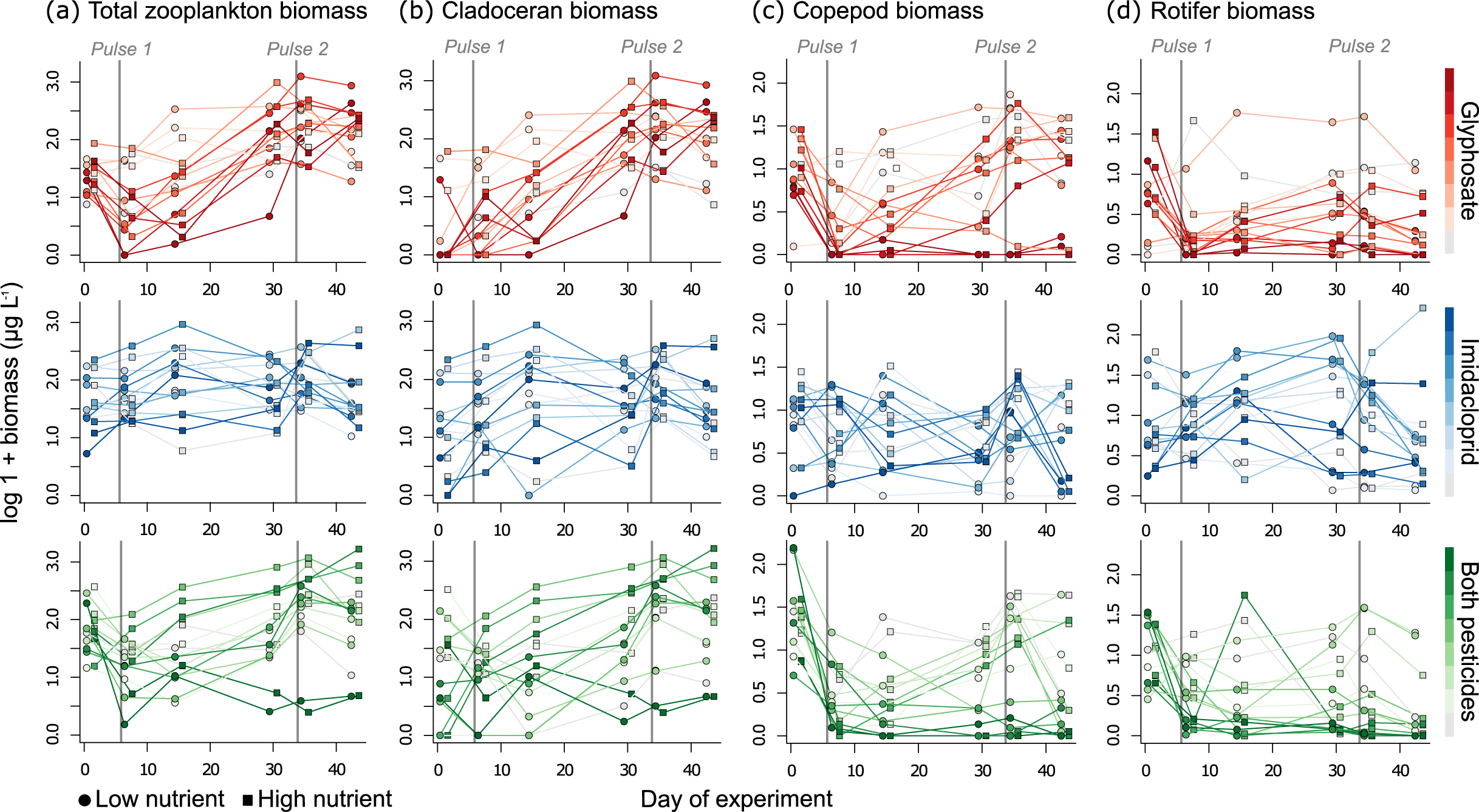
Temporal dynamics of (a) total zooplankton, (b) cladoceran, (c) copepod, and (d) rotifer community biomass over the course of the experiment. Symbols correspond to low- and high-nutrient treatments; colour type and saturation indicate the nature and dose of pesticide treatment, respectively. A small horizontal offset between low- and high-nutrient ponds was used to facilitate visualisation.

**FIGURE 3.**
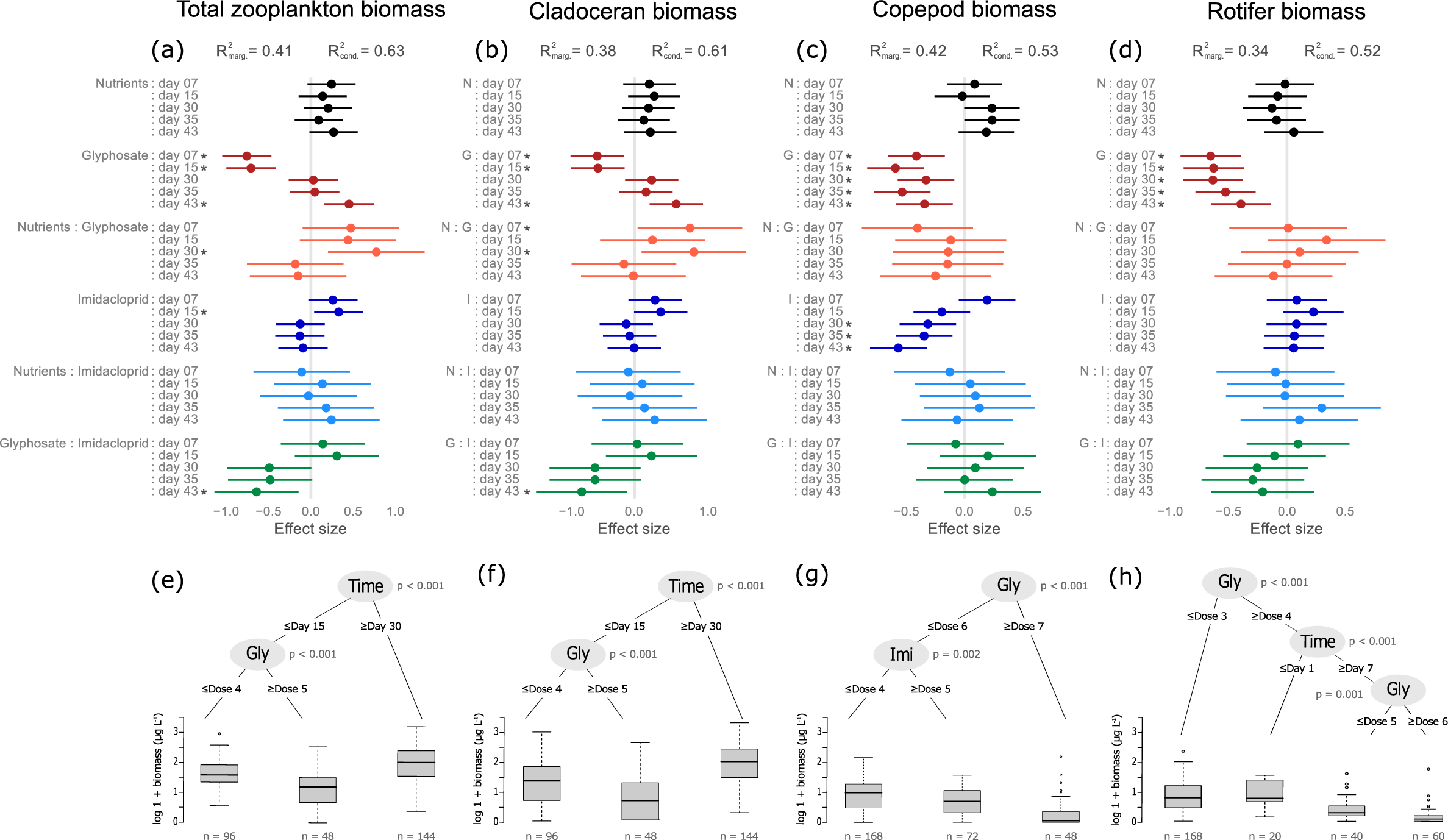
Effects of agrochemical treatments on zooplankton biomass. (a-d) Forest plots illustrating the time-dependent effects of nutrients, glyphosate and imidacloprid and their interactions on (a) total zooplankton, (b) cladoceran, (c) copepod, and (d) rotifer community biomass. In forest plots, each point represents an effect size quantified as the parameter estimates of linear mixed effect models (LMMs). Error bars represent 95% confidence intervals, with effects being significant when there is no overlap with zero; asterisks highlight significant effects. Table S3 provides a full list of model coefficients. (e-h) Univariate regression tree (URT) models identifying thresholds (agrochemicals or day of experiment) of response in (e) total zooplankton, (f) cladoceran, (g) copepod, and (h) rotifer community biomass.

One day after the first pulse of pesticides (day 7), glyphosate triggered a rapid decline in total zooplankton biomass (Figures 2a, 3a). While the application of glyphosate (alone or in combination) clearly triggered the collapse of copepods and rotifers (Figure 2c-d), no such collapse occurred in cladocerans (Figure 2b). At low and moderate glyphosate concentrations, copepod biomass remained relatively low but partly recovered over time; while higher doses of glyphosate resulted in a long-lasting collapse, especially in immature copepods (Figures 2c, S2c-e). Rotifers entirely collapsed in ponds exposed to glyphosate (alone or in combination) and failed to recover (Figure 2d). For cladocerans, the negative effect of glyphosate was transient, only visible on days 7 and 15 (Figures 2b, 3b). Cladocerans then markedly increased over time, with no apparent decline even after the second pesticide pulse. By day 43, a strong, positive effect of glyphosate was detected (Figure 3b), with increasing cladoceran biomass along the glyphosate gradient (Figures 2b, S2b). This was primarily driven by increases in *Alona sp*., *Chydorus sphaericus*, and *Scapholeberis mucronata* (hereafter referred by genus only; Figure S1a, d, g). Because cladocerans constituted a large proportion of zooplankton biomass, the time-dependent effect of glyphosate on Cladocera, shifting from a negative to a positive influence over time, was also visible for total biomass (Figures 2a, 3a). Importantly, total zooplankton biomass showed clear recovery from pulse 1, and no apparent decline after pulse 2.

Imidacloprid had a clear negative effect on copepod biomass over time (Figures 3c, S2c-e), especially in copepodites after pulse 2 (Figure S1l). However, imidacloprid did not affect cladoceran, rotifer, or total zooplankton biomass. A weak positive effect was detected on total zooplankton on day 15, likely due to marginally significant positive effects on rotifers and cladocerans (Figure 3a-d).

Overall, the joint presence of glyphosate and imidacloprid resulted in responses akin to those observed in ponds treated with glyphosate alone (Figure 2). Temporal increases in *Alona, Chydorus*, and *Scapholeberis* biomass were observed across ponds treated with any pesticide (Figure S1a, d, g), indicating a (co-)tolerance to both pesticides. One notable exception were ponds exposed to the highest dose of both pesticides, where cladocerans, and thus zooplankton biomass, declined dramatically and never recovered (Figure 2a-b), reflecting strong pesticide interaction. A LMM revealed that pesticide interactions on total zooplankton and cladoceran biomass were, however, only significant on day 43 (Figure 3a-b). This result constitutes the only evidence of strong agrochemical interaction in our study; only few weak interactions were found otherwise.

Overall, glyphosate was the strongest driver of zooplankton biomass (Figure 3). A URT identified threshold concentrations of glyphosate causing substantial biomass decreases in rotifers (dose 4 = 0.3 mg/L), cladocerans (dose 5 = 0.7 mg/L; transient negative effect only), and copepods (dose 7 = 5.5 mg/L; Figure 3f-h). Dose 5 (0.7 mg/L) was also the threshold exposure at which total biomass started to decrease (transient effect; Figure 3e). A breaking point was also identified for imidacloprid, with concentrations ≥3µg/L (dose 5) triggering declines in copepods biomass.

In a separate analysis, a LMM indicated that chl-*a* concentration was also a driver of zooplankton biomass, but with limited predictive power. Chl-*a* enhanced total zooplankton and cladoceran biomass, but had little to no effect on copepods and rotifers (Figure S3; see Table S4 for further detail). By the end of the experiment, ponds treated with glyphosate or both pesticides had the highest biomass of algae and zooplankton, except for the combination of the highest pesticide doses (Figure S3). Given that chl-*a* was (i) highly correlated with glyphosate and TP after pulse 1 and (ii) a weaker driver of zooplankton biomass than agrochemicals, chl-*a* was excluded from the set of predictors (see Methods).

### 3.2 Effects of agrochemicals on community structure

A total of 24 zooplankton taxa were identified in this experiment, reflecting the complexity of our semi-natural pond communities.

#### Community composition

A series of six ordinations (PCA) uncovered a clear effect of time and treatment on zooplankton community composition. On day 1, communities were similar across ponds (Figure 4a). Shortly after the first pesticide pulse, communities diverged in composition, with ponds exposed to glyphosate versus imidacloprid being diametrically opposed in the ordinations of days 7 and 15 (Figure 4b-c). Community assemblages remained different across ponds until the end of the experiment (Figure 4d-f). Using PC1 scores as a proxy for community composition, both the LMM and URT indicated that glyphosate and time were the strongest drivers of compositional changes (Figure 4g-h; Table S5), with the greatest compositional shift occurring at ≥2 mg/L of glyphosate (dose 6).

**FIGURE 4.**
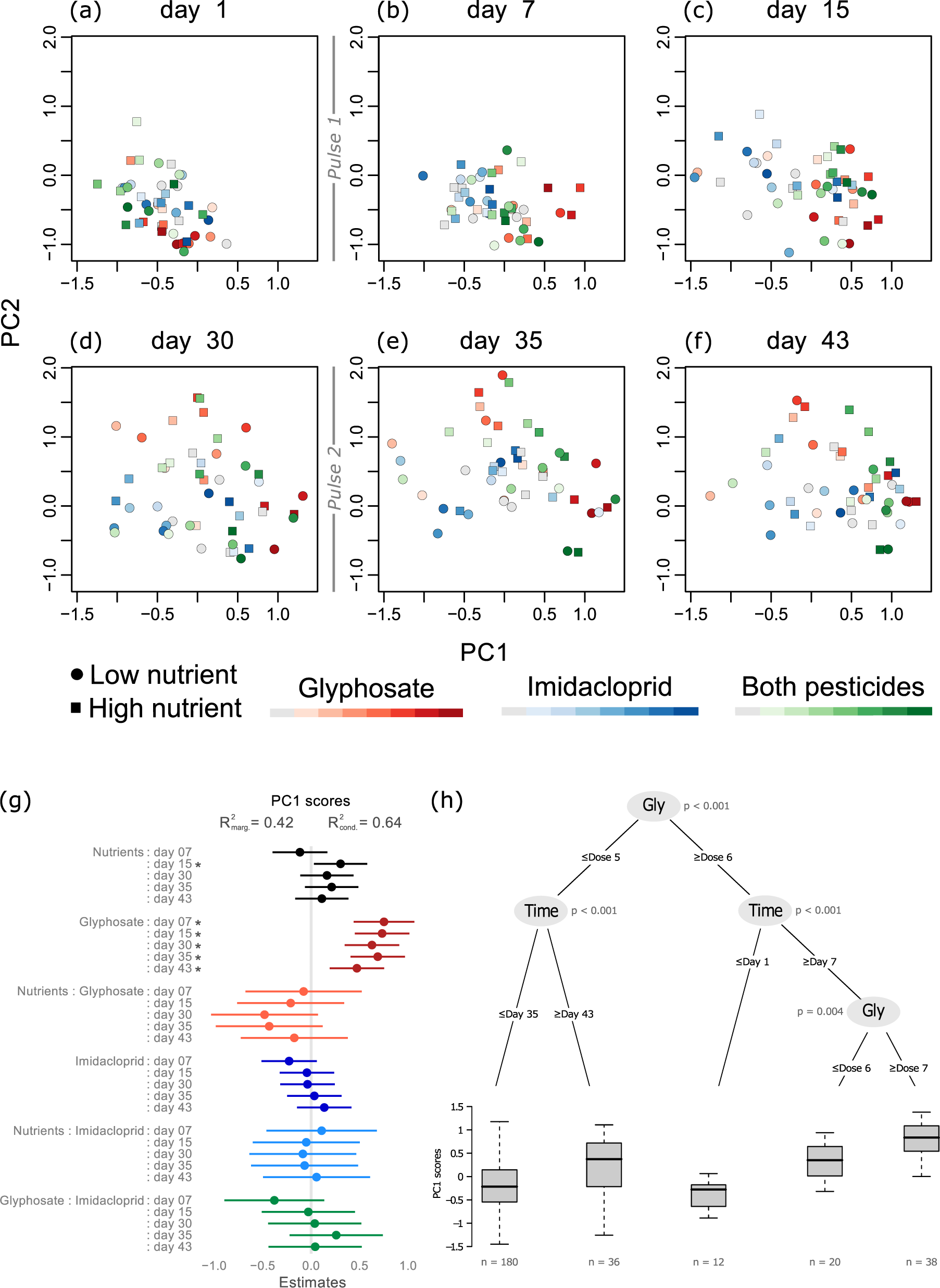
Temporal variation in zooplankton community composition across ponds and agrochemical treatments. (a-f) Principal component analysis (PCA) illustrates changes in community composition across sampling days. (g) Forest plot indicating the time-dependent effects of agrochemicals and their interactions on changes in community composition (measured as PC scores of the first axis shown in a-f; PC1). Effect sizes are quantified as the parameter estimate of LMMs; however, direction of effects should not be interpreted. See Table S5 for a full list of model coefficients. (h) URT model identifying thresholds (agrochemicals or day of experiment) of response in community composition (measured as PC1) over the course of the experiment.

#### Responsive taxa

PRCs illustrated compositional divergence between treated and control ponds (deviations of coloured lines from the gray horizontal line, where Y=0), while indicating taxon-level responses (right Y axis reflecting increases or decreases in most responsive taxa). The first pulse of glyphosate triggered a decline in several species (Figure 5a-b). By day 30, communities faced with low to moderate treatments showed an increase in *Scapholeberis* and other cladocerans. Ponds exposed to high glyphosate doses lost most species present in less contaminated ponds (e.g., copepods, rotifers; Figures 5a-b), being primarily composed of *Alona* (distinctly high density) and *Chydorus* (Figure S1a, d). Imidacloprid also induced compositional changes (Figure 5c-d). However, apart from a greater representation of some rotifers and an increased presence of *Scapholeberis*, no clear pattern of species turnover emerged along the insecticide gradient; in contrast to glyphosate.

**FIGURE 5.**
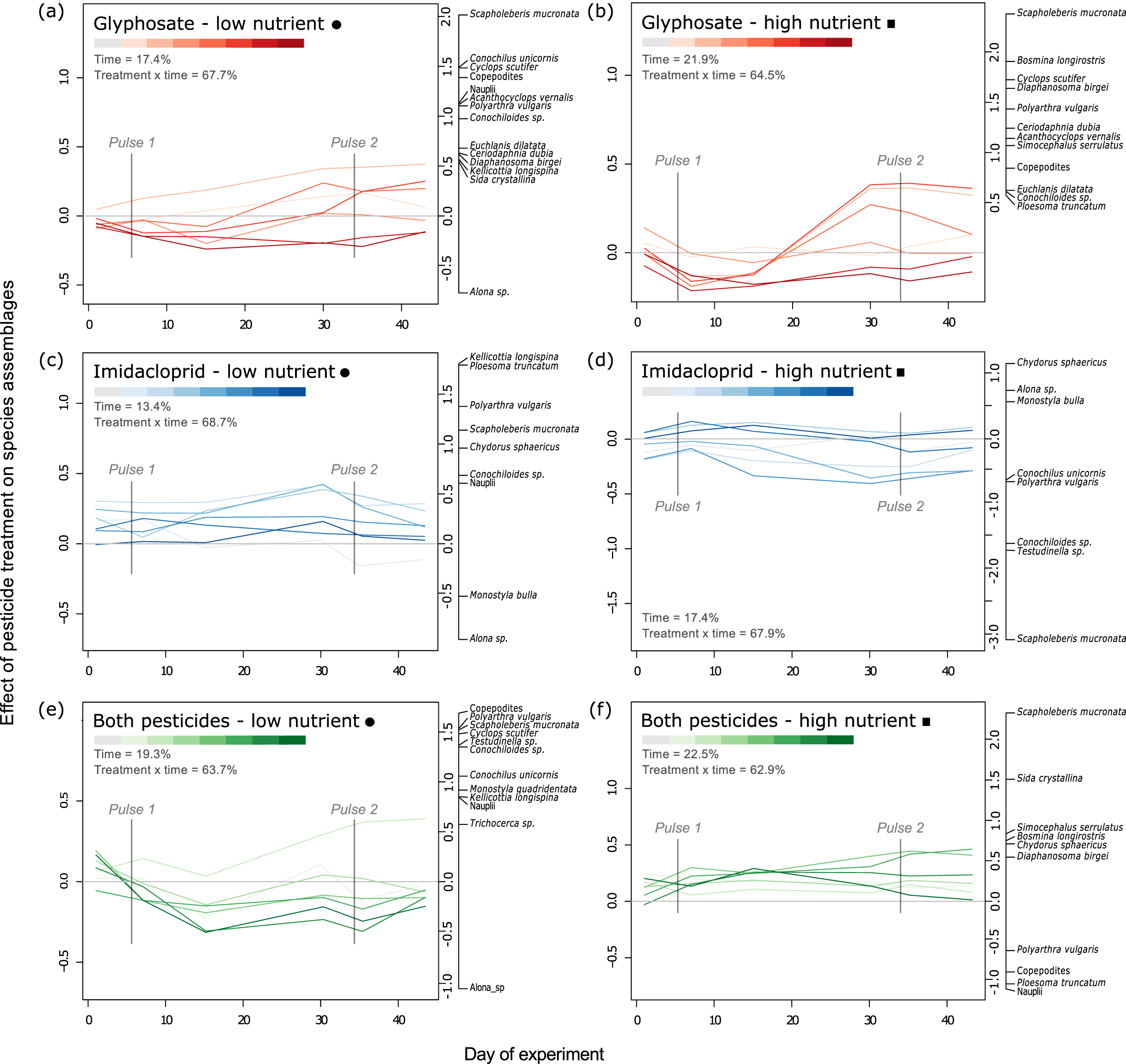
Temporal variation in zooplankton community assemblages illustrated by principal response curves (PRCs). PRCs graphically represent how community composition diverge between control ponds and ponds exposed to (a-b) glyphosate, (c-d), imidacloprid, and (e-f) both pesticides, under low- (a, c, e) or high- (b, d, f) nutrient conditions. The left Y axis reflects deviations in community composition (curves) over time, whereas the right Y axis indicate the relative contribution of species to community compositional changes (species scores).

Communities confronted with both pesticides showed clear compositional shifts, with stronger turnover at higher doses (Figure 5e-f). Certain taxa responded similarly when faced with glyphosate alone and both pesticides (Figure 5a-b versus e-f); e.g., declines in rotifers (Figure S1). In low-nutrient ponds, assemblages deviated distinctly from controls after each pulse, with *Alona* becoming dominant whilst rotifers and copepods decreased in density (Figure 5e). In high-nutrient ponds, communities were characterised by an increase in cladocerans, especially *Scapholeberis*, and a clear loss of rotifers and immature copepods (Figure 5f).

The interaction between time and nutrient enrichment alone explained relatively little variation in species assemblages (36%; Figure S4), indicating that pesticide treatment was a stronger driver of species turnover than nutrient enrichment.

#### Diversity metrics

Pesticide type and dose affected zooplankton taxon richness and, thus, alpha diversity (Shannon index; Figure S5a-f), but had no apparent effect on community evenness (Figure S5g-i). LMMs using richness as a response variable provided a better fit than models using alpha diversity for all of total zooplankton, crustacean, and rotifer communities (Figures 6a-c versus S6a; Tables S6-S7). Thus, we primarily focus on effects of agrochemicals on richness, while reporting analogous but weaker effects on other measures of diversity in the Supplementary Material.

**FIGURE 6.**
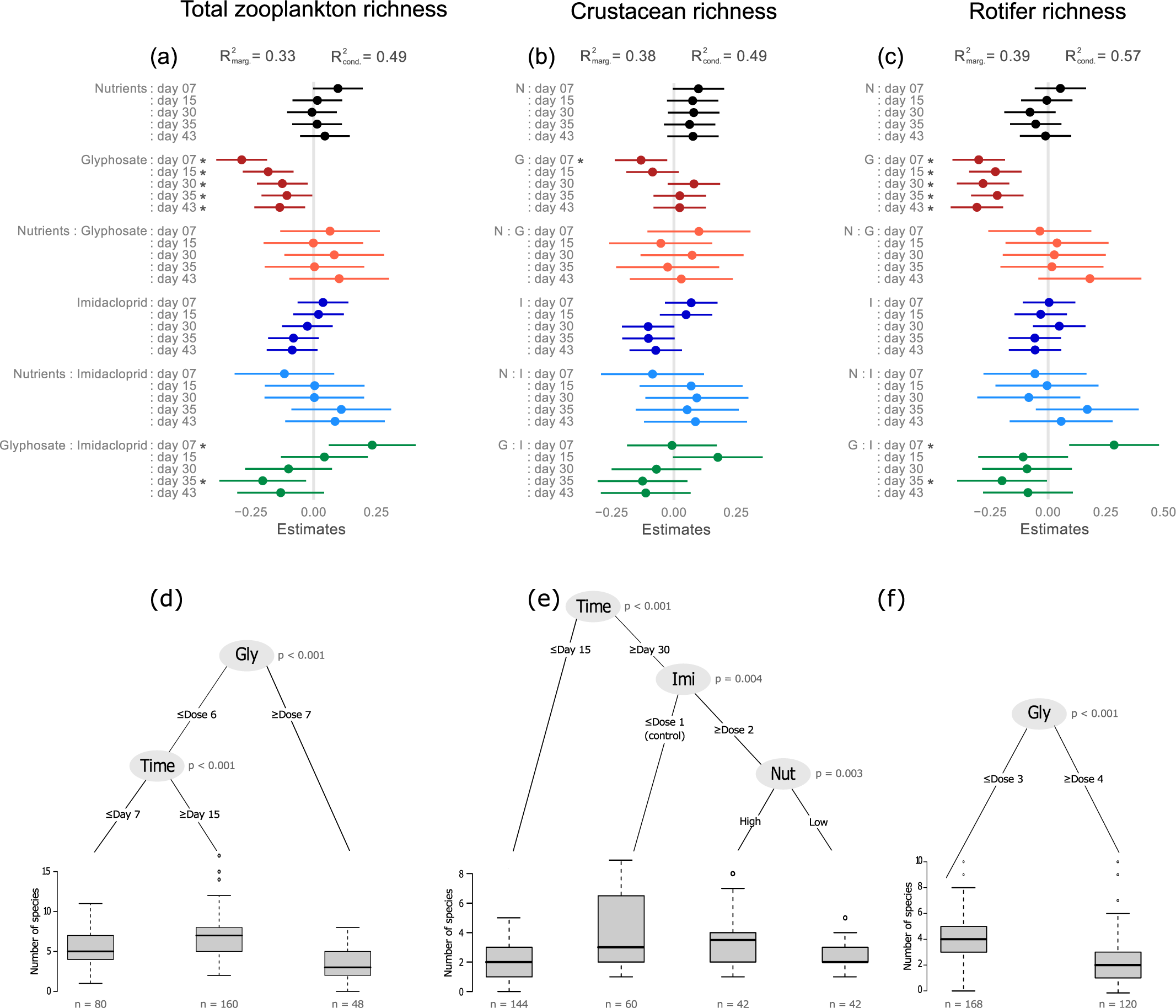
Effects of agrochemical treatments on the number of zooplankton taxa. (a-c) Forest plots illustrating the time-dependent effects of agrochemicals and their interactions on (a) total zooplankton, (b) crustacean, and (c) rotifer richness over time. Effect sizes are quantified as the parameter estimate of LMMs; see Table S6 for a full list of model coefficients. (d-f) URT models identifying thresholds (agrochemicals or day of experiment) of response in (d) total zooplankton, (e) crustacean, and (f) rotifer richness.

**FIGURE 7.**
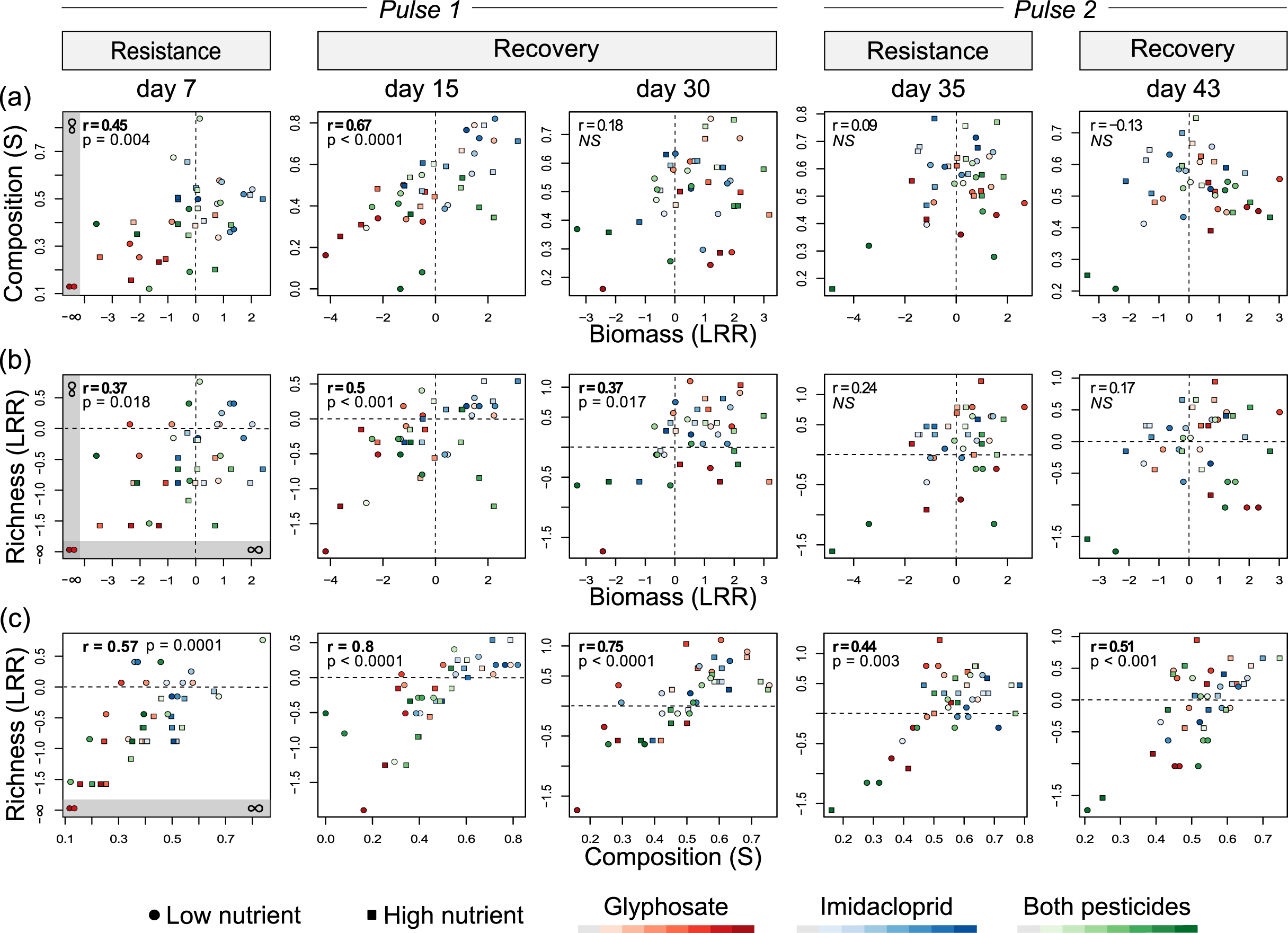
Relationships between the resistance or recovery of zooplankton community properties over time: a) composition versus biomass; b) richness versus biomass; c) richness versus composition. Resistance measures are estimated on days 7 (pulse 1) and 35 (pulse 2), whereas recovery measures are estimated on days 15, 30 (i.e., 9 and 24 days after pulse 1), and 43 (9 days after pulse 2). Spearman rank correlation coefficients are indicated for each bivariate relationship; significant coefficients (*P* < 0.05) are highlighted in bold. Resistance and recovery measures of biomass and richness are expressed as log response ratio (LRR) between treated and control communities: 0 = maximum resistance or recovery, as indicated via dash lines; <0 = low resistance via alteration/underperformance or incomplete recovery; >0 low resistance via overperformance/stimulation or recovery via overcompensation/stimulation. For composition, measures are expressed as a similarity index (S) between treated and (nutrient-specific) control communities: 0 = low; 1 = maximum resistance/recovery. Note that in instances where entire zooplankton communities momentarily collapsed (i.e., absence of individuals in 2 low-nutrient high-glyphosate ponds on day 7), LRRs based on null values of biomass and richness = -∞; such measures are graphically represented but excluded from correlation coefficients.

Glyphosate strongly affected zooplankton richness (Figure 6a-c), especially in rotifers. Unlike its effect on biomass, glyphosate maintained an adverse effect on community richness throughout the experiment (Figures 6a versus 3a). Declines in richness were also stronger with glyphosate doses, applied alone or in combination (Figure S5a-c); however, crustaceans exposed to low doses of glyphosate alone (not in the presence of imidacloprid) increased their richness over time (Figure S5a-b). A URT determined that ≥0.3 mg/L (dose 4) of glyphosate resulted in a two-fold decrease in rotifer richness (Figure 6f); an even lower breaking point (0.1 mg/L glyphsoate) was identified for alpha diversity (Figure S6b). Imidacloprid (≥0.15 µg/L) also reduced crustacean richness as a result of copepod sensitivity, but only under low-nutrient conditions. Imidacloprid was the only driver of crustacean diversity loss (Figure S6b).

### 3.3 Resistance and recovery of community properties

For the first pesticide pulse, community resistance and recovery were lower with increasing pesticide treatments in all of the community properties (Figure S7). Resistance and recovery were generally stronger for pulse 2 relative to pulse 1 (LRRs closer to 0; composition similarity closer to 1). For biomass only, the long-term recovery was stronger with increasing pesticide doses, except for the highest combination of both pesticides (Figure S7a). By the end of the experiment, while pond richness and composition had partially recovered, biomass had exceeded full recovery (surpassing biomass levels in control ponds), showing clear signs of stimulation by glyphosate. Given the strong correlation between richness and other measures of diversity, results for the latter are reported in the Supplementary Material.

Aside from reflecting a relatively smaller effect of the second pulse on zooplankton communities, the analysis of resistance and recovery within community properties was inconclusive (Figure S9). In particular, none of the community properties showed a correlated resistance to both pulses (Figure S9a).

The examination of relationships among community properties revealed that the responses in biomass and composition were positively correlated upon and after pulse 1 (days 7 and 15; Figure 7a); by day 30, however, biomass markedly recovered, unlike composition. Similarly, richness and biomass responses were correlated for pulse 1, but not pulse 2 (Figure 7b); reflecting the greater recovery of biomass relative to richness over time. Composition and richness remained tightly coupled over the experiment, indicating that compositional shifts were primarily owed to changes in the number of taxa (Figure 7c).

At the end of the experiment (day 43), the relationship between biomass and composition had become visibly negative (Figure 7a); yet, no significant correlation was detected due to the absence of biomass recovery in the two pond communities faced with strong pesticide interaction at the highest combination of treatments (isolated in the lower left quadrant). Indeed, the negative correlation becomes significant (r=-0.35; *P*=0.05) when these two ponds are removed, indicating a potential trade-off between biomass and composition recovery across all other 46 ponds.

## 4. Discussion

Combining approaches from community ecology and ecotoxicology in the context of agricultural pollution, we quantified the effects of three widespread synthetic agrochemicals on zooplankton biomass and community structure. Of all possible single and interactive effects, we found that glyphosate, applied alone or in combination, was the most influential driver of community-wide biomass, composition, and diversity metrics, with a marked time-dependent effect on biomass. Although imidacloprid distinctly impaired copepods, the insecticide did not affect zooplankton community-wide biomass. Our results indicate that community properties respond rapidly to glyphosate exposure but may not recover equally over time. Importantly, at the community scale, zooplankton biomass showed little resistance to the first pulse of glyphosate, but rapidly recovered and even increased as a function of the glyphosate dose received; in ponds exposed to higher concentrations of glyphosate (alone or with imidacloprid), however, the number of taxa declined and never recovered over the experiment. We found that some cladocerans can be highly tolerant to pesticide contamination, and their biomass can compensate for the decline of more sensitive taxa; this sorting process conferred greater resistance to zooplankton communities during the second pulse. Below, we position the pesticide concentrations inducing biotic responses within the ecotoxicological literature (reviewed in Table S1), and discuss our findings with regards to underlying community processes, as well as implications for freshwater zooplankton in agricultural areas.

### 4.1 Community responses to agrochemicals

Zooplankton biomass and structural (composition, diversity) responses varied with agrochemical treatment type and severity, and among major taxonomic units (cladocerans, copepods, rotifers). Unlike other studies (Alexander et al., 2016; Baker et al., 2016; Geyer et al., 2016), nutrient enrichment, alone or combined with pesticide contamination, had surprisingly little effect on zooplankton in our ponds. Of the few nutrient-related signals detected, the positive interactive effect with glyphosate on cladoceran (and thus total zooplankton) biomass after the first pesticide pulse was the strongest, indicating that nutrients partly contributed to the subsequent cladoceran proliferation and overall zooplankton community recovery.

Although imidacloprid exerted milder effects on zooplankton communities as compared to glyphosate, the insecticide distinctly impaired copepods. Given that only two copepod species were identified in our ponds, this result should not be generalised across Copepoda. Copepod biomass declined at doses ≥3 µg/L, in agreement with other mesocosm-based assessments reporting harmful effects at concentrations between 3 and 4 µg/L (Chará-Serna et al., 2019; Schrama et al., 2017; Sumon et al., 2018). A compelling case study led by Yamamuro et al. (2019) showed that decadal increases in the use of neonicotinoids caused the collapse of freshwater zooplankton, and in turn, of fisheries yield; notably, the once dominant and most sensitive taxon of the zooplankton community was a copepod, whose biomass decline coincided with the introduction of imidacloprid in 1993. Unlike copepods, cladocerans and rotifers appeared to be tolerant overall to imidacloprid in our experiment. Yet, adverse effects of imidacloprid have been documented at similar or even lower levels for a wide array of aquatic invertebrates (Morrissey et al., 2015; Raby et al., 2018; Van Dijk et al., 2013), including some cladocerans and rotifers found in our ponds (e.g., *Polyarthra*; Sumon et al., 2018; Table S1). No clear patterns of species sorting emerged along the imidacloprid gradient; however, the decline in copepods likely benefitted cladocerans and rotifers after the first pulse, possibly as a result of relaxed competition. In contrast to findings reported by Chará-Serna et al. (2019), the sole presence of imidacloprid in our ponds decreased the number of crustacean taxa; however, nutrient enrichment neutralised this effect, as observed in previous studies (Alexander et al., 2013; 2016).

The presence of glyphosate in ponds resulted in differential short- and long-lasting effects across major zooplankton groups. Indeed, the first pulse triggered immediate declines in copepods and rotifers but not cladocerans, while subsequent biomass increases were only observed in cladocerans. This pattern was clearly visible in ponds contaminated with the herbicide alone and in conjunction with the insecticide, highlighting the relatively strong influence of glyphosate in our experiment, but more importantly, the co-tolerance of some cladocerans to pesticides. However, the prolonged collapse of all major groups, and thus of overall zooplankton, in ponds treated with the combination of the highest doses of pesticides reveals evidence of negative interactive effects of glyphosate and imidacloprid on zooplankton, which has thus far not been documented, to our knowledge.

The general patterns of species turnover in communities exposed to glyphosate (alone or in combination) highlighted a stark contrast between sensitive rotifer (long-lasting collapse at concentrations ≥0.3 mg/L) and copepod zooplankton (partial recovery over time, but none ≥5.5 mg/L), and highly tolerant cladocerans, primarily belonging to *Alona, Chydorus*, and *Scapholeberis* (still thriving at 15mg/L of glyphosate, even in the presence of imidacloprid). Information on rotifers is relatively limited (Table S1), but our results are consistent with other studies of glyphosate toxicity on copepods, with notably greater sensitivity found in immature stages as compared to adults (Lim et al., 2019). While compelling evidence has accrued for acute and chronic toxicity in cladocerans at similar concentrations of glyphosate (Table S1; e.g., ranges of LC_50_ for *Daphnia* and *Ceriodaphnia*: 0.45–10.6 and 4.8–6.05 mg/L, respectively; Cuhra et al., 2013; Tsui & Chu, 2003), such observations mostly rely on single-species cultures in a laboratory setting, with few to no studies using the most tolerant species in our ponds. Several multispecies assessments recorded only minor or transient community-wide density effects of glyphosate (Baker et al., 2016; Gutierrez et al., 2017; Lu et al., 2020; Vera et al., 2012), and concluded that the herbicide was unlikely to cause the collapse of entire zooplankton communities under normal-use circumstances (e.g., glyphosate level up to ∼2.25 mg/L in water, as per Relyea, 2005; 2006). In this regard, our results also suggest that, on a longer-term basis, glyphosate may have limited adverse effects on total zooplankton biomass. Nonetheless, the sensitivity and long-lasting collapse of rotifers may have important implication for food web processes (Arndt, 1993; Miracle et al., 2007), warranting further investigation.

The rapid and marked proliferation of tolerant cladocerans subsequent to the first application of glyphosate was likely attributable to a stimulatory, bottom-up effect induced by the nutrient content of glyphosate. Glyphosate acid contains 18.3% P, implying that its presence in water represents an additional source of P that may be used by microbial and algal communities, either in the form of glyphosate or degraded products (Brock et al., 2019; Hove-Jensen et al., 2014; Wang et al., 2016; 2017). Other studies have reported increases in phytoplankton in the presence of glyphosate, attributing this fertilising effect to glyphosate-derived P (Forlani et al., 2008; Harris & Smith, 2016; Pérez et al., 2007; Saxton et al., 2011). The parallel study by Fugère et al. (2020) showed that glyphosate led to dose-dependent increases in TP and chl-*a* concentrations in our ponds, presumably as a result of phytoplankton P-limitation (initial N:P ratio ∼33). Taken in context with our results, it appears that glyphosate-mediated increases in algal resources enhanced the growth of tolerant cladocerans. While positive (indirect) effects of glyphosate on cladoceran and overall zooplankton biomass were visible at all treatment concentrations by the end of our experiment, the highest biomass levels were recorded in the most contaminated ponds, demonstrating that the dose-dependent fertilising effect of glyphosate on phytoplankton can transfer to higher trophic levels. Though less documented than toxic effects, bottom-up effects of glyphosate on zooplankton have been previously observed (Vera et al., 2012). Overall, our results reveal a two-step effect of glyphosate on zooplankton biomass: negative in the short-term due to the loss of sensitive taxa, but positive over longer timescales owing to the increased growth of tolerant taxa. This positive effect will however be conditional upon the presence of at least one or a few tolerant taxa in the community, a condition that may not always be satisfied.

As a result of differential sensitivity across taxa, the rise in (co-)tolerant cladoceran zooplankton permitted the recovery of community biomass in ponds treated with glyphosate or both pesticides; this sorting process also prompted greater resistance during the second pulse. This result is consistent with the concept of stress- or pollution-induced community tolerance (Bérard & Benninghoff, 2001; Tlili et al., 2016; Vinebrook et al., 2004), whereby initial exposure to stress may eliminate sensitive taxa from a community, leading to increased community tolerance upon subsequent exposure. Indeed, species sorting in favour of tolerant cladocerans induced by the first pulse of glyphosate in our ponds likely conferred community tolerance (and thus greater biomass resistance), allowing maintenance of zooplankton biomass following the second pulse. This result constitutes evidence of pollution-induced community tolerance in zooplankton faced with pesticide contamination, providing a rare example of such tolerance in metazoans, given that most ecotoxicological studies of pollution-induced community tolerance focus on microbial, algal, or periphytic communities (Blanck, 2002; Boivin et al., 2002; Tlili et al., 2016).

We also found that community-wide biomass showed greater recovery than composition and diversity, as have previous studies of functional and structural responses to environmental disturbances (Hillebrand & Kunze, 2020; Hoover et al., 2014). In fact, the recovery and increase of biomass after exposure to glyphosate (alone or with imidacloprid) was achieved through species sorting, all while community richness and (alpha) diversity declined in the more contaminated ponds. The striking contrast in temporal patterns of biomass versus richness, whereby effects of glyphosate were time-dependent for the former but remained negative for the latter (Figures 3a vs 6a), have clear implications for biodiversity loss and ecosystem functioning in freshwaters; that is, even when total zooplankton biomass appears unaffected. When excluding the two pond communities that failed to recover in biomass (i.e., those faced with strong pesticide interaction at the highest treatment combination), we also found a significant negative relationship between recovery of community biomass and composition, highlighting the contrasting response of these properties over time and pointing to a potential trade-off between the recovery of biomass and community structure. In sum, our study indicates that the long-term effect of glyphosate contamination on zooplankton varies among community properties, with increasing concentrations triggering clear compositional shifts, species loss, but greater biomass production in the remaining tolerant taxa.

### 4.2 Implications and concluding remarks

Comprising a total of 24 zooplankton taxa representative of the local species pool (Thompson et al., 2015), our 48 semi-natural pond communities revealed complex processes that may only be observable under realistic conditions: differential patterns of sensitivity and recovery, glyphosate-mediated bottom-up effects, and pesticide interactions. Together, these results integrate ecotoxicological and ecological approaches to assess effects of synthetic contaminants on biota, while also providing insight into the study of community stability and multiple stressors in the context of agrochemical pollution. Our study responds to calls for more complex multispecies testing approaches in ecotoxicology and better inform risk assessments and conservation practices (Clements & Rohr, 2009; Gessner & Tlili, 2016; Relyea & Hoverman, 2006).

Although community biomass may be resilient to severe pesticide contamination, species loss and compositional shifts in favour of a few distinct tolerant taxa can have implications for food web processes and ecosystem stability in agricultural areas (Frank & Tooker, 2020; Pennekamp et al., 2018). Losing taxon-specific functions while enhancing those provided by tolerant taxa could destabilise ecosystems (Arnoldi et al., 2019). In environments prone to glyphosate pollution, the proliferation of herbivorous cladocerans at the expense of omnivorous or carnivorous cyclopoids or less effective filter-feeding rotifers could modulate trophic interactions and top down pressure (Sommer et al., 2001; 2003). Another potential consequence of glyphosate is nutrient enrichment cascading up the food chain. While often overlooked as a source of anthropogenic P in agricultural landscapes, glyphosate-derived P inputs are now comparable in magnitude to other past P-sources (e.g., detergents) that once required legislation (Hébert et al., 2019), and its excessive usage warrants more attention in watershed management.

In this experiment, concentrations at which pesticides caused biotic effects differed among zooplankton community properties, with most response thresholds falling within the range of previously recorded levels in agricultural water bodies. Though pervasive in surface waters, imidacloprid and glyphosate concentrations are highly variable, in part owing to variability in land use intensity and biodegradation potential (Giroux, 2019; Hladik et al., 2014; 2018; Medalie et al., 2020). Globally, imidacloprid and glyphosate concentrations can range between <0.01–320 µg/L and <0.001–5.2 mg/L, respectively (Annett et al., 2014; Hénault-Éthier et al., 2017; Morrissey et al., 2015; Struger et al., 2008; 2017); but levels on the order of ng/L are common for both pesticides (Montiel-Léon et al., 2019). In our ponds, glyphosate adversely affected overall zooplankton biomass at 0.7 mg/L (≥0.3 mg/L in rotifers), but concentrations ≥0.04 mg/L were sufficient to enhance total biomass over time; and alterations of community structure were observed at 0.1 mg/L. For imidacloprid, concentrations ≥3 µg/L affected copepod biomass and ≥0.15 µg/L reduced crustacean richness and diversity. While some of these threshold exposure concentrations may be on the high end of environmentally relevant glyphosate concentrations, especially for glyphosate, most effects were observed under the benchmark of 2.25 mg/L that is often used as the worst-case scenario in ecotoxicology (Geyer et al., 2016; Relyea, 2005; 2006).

Most crucially, our study demonstrates that community properties could be affected at pesticide levels below common North American water quality guidelines. This is especially the case for glyphosate benchmarks for the protection of aquatic life in Canada (0.8 and 28 mg/L; CCME 2007; 2012) and the United States (for freshwater invertebrates: 26.6 and 49.9 mg/L; EPA 2019). Although Canadian criteria for glyphosate remain more conservative as compared to those of the United States, these criteria appear too permissive to ensure protection. In light of the global expansion in glyphosate use, we believe that such guidelines should be re-evaluated to meet conservation goals in agricultural areas.

## Acknowledgements

We are thankful to Michelle Gros, Katherine Velghe, Marilyne Robidoux, and Marco Castro for their assistance with laboratory work; David Maneli, Charles Normandin, Alexandre Arkilanian, and Tara Jagadeesh for their help with field work; Capucine Lechartre and Olympe Durrenberger for their assistance with the literature review. MPH was supported by the Natural Sciences and Engineering Research Council (NSERC), Fonds de Recherche du Québec Nature-Technologies (FRQNT), and NSERC-CREATE via the GRIL. VF was supported by NSERC; NBC was supported by FRQNT and NSERC-CREATE via the GRIL. The LEAP facility was funded by the Canadian Foundation for Innovation and Liber Ero Chair in Biodiversity Conservation attributed to AG. This work was also supported by a National Geographic grant for early-career researchers awarded to MPH and NSERC Discovery Grants attributed to BEB, AG, and GFF.

